# Atrial fibrillation associated common risk variants in SYNE2 lead to lower expression of nesprin-2α1 and increased nuclear stiffness

**DOI:** 10.1101/708057

**Authors:** Nana Liu, Jeffrey Hsu, Gautam Mahajan, Han Sun, John Barnard, David R. Van Wagoner, Chandrasekhar R. Kothapalli, Mina K. Chung, Jonathan D. Smith

**Author notes:** Address correspondence to: Dr. Jonathan D. Smith, Box NC10, Cleveland Clinic, 9500 Euclid Avenue, Cleveland, OH 44195.

## Abstract

**Rationale:** Atrial fibrillation (AF) genome-wide association studies (GWAS) identified significant associations for rs1152591 and linked variants in the SYNE2 gene encoding the nesprin-2 protein that connects the nuclear membrane with the cytoskeleton

**Objective:** Determine the effects of the AF-associated rs1152591 and rs1152595, two linked intronic single nucleotide polymorphisms (SNPs), on SYNE2 expression and investigate the mechanisms for their association with AF.

**Methods and Results:** RNA sequencing of human left atrial appendage (LAA) tissues indicated that rs1152591 and rs1152595 were significantly associated with the expressions of SYNE2α1, a short mRNA isoform, without an effect on the expression of the full-length SYNE2 mRNA. SYNE2α1 mRNA uses an alternative transcription start site and encodes an N-terminal deleted 62 kDa nesprin-2α1 isoform, which can act as a dominant-negative on nuclear-cytoskeleton connectivity. Western blot and qPCR assays confirmed that AF risk alleles of both SNPs were associated with lower expression of nesprin-2α1 in human LAA tissues. Reporter gene transfections demonstrated that the risk vs. reference alleles of rs1152591 and rs1152595 had decreased enhancer activity. SYNE2 siRNA knockdown (KD) or nesprin-2α1 overexpression studies in human stem cell-derived induced cardiomyocytes (iCMs) resulted in ~12.5 % increases in the nuclear area compared to controls (p<0.001). Atomic force microscopy demonstrated that SYNE2 KD or nesprin-2α1 overexpression led to 57.5% or 33.2% decreases, respectively, in nuclear stiffness compared to controls (p< 0.0001).

**Conclusions:** AF-associated SNPs rs1152591 and rs1152595 downregulate the expression of SYNE2α1, increasing nuclear-cytoskeletal connectivity and nuclear stiffness. The resulting increase in mechanical stress may play a role in the development of AF.

## INTRODUCTION

Atrial fibrillation (AF), the most common clinically significant arrhythmia, is characterized by rapid and irregular activation in the atria without discrete P waves on the surface electrocardiogram (ECG). It is estimated that there are 5.2 million AF patients in the United States, and the number is expected to increase to 12.1 million over the next 1-2 decades^1^. This disorder is a major cause of morbidity, mortality, and health care expenditure. Two AF genome-wide association studies (GWAS) in 2012 and 2017 identified the single nucleotide polymorphisms (SNP) rs1152591, located in an intron of SYNE2 on chromosome 14q23, as highly significant ^2, 3^. Our prior RNA sequencing of human left atrial appendages (LAA) from a biracial cohort of 265 subjects found that rs1152591 is associated with SYNE2 gene expression, thus acting as a cis-expression quantitative trait locus (cis-eQTL) for SYNE2, with the risk allele associated with lower SYNE2 expression ^4^.

Nesprin-1/2 are two related outer nuclear membrane-spanning proteins, encoded by the SYNE1/2 genes. Alternative promoters, RNA splicing, and termination give rise to multiple isoforms of nesprin-1/2. The longest isoform of nesprin-2, referred to as the giant isoform, contains three different functional domains: the C-terminal transmembrane KASH (Klarsicht-ANC-Syne-homology) domain, which anchors within nuclear membrane, multiple spectrin-repeat (SR) rod domains, and the N terminal paired calponin-homology (CH) domain connecting with cytoskeletal actin. Due to lack of the CH domain, nesprin-2 short isoforms localize mainly to the nuclear membrane without binding to cytoskeletal actin. Nesprin-2 giant binds to SUN proteins (Sad1p/UNC-84) between the outer and inner nuclear membranes, which in turn span the inner membrane to bind lamin A/C underneath the inner nuclear membrane and compose the LINC (Linker of Nucleoskeleton and Cytoskeleton) complex, which mechanically links the nucleoskeleton to the cytoskeleton ^5–7^. The LINC complex has diverse functions, including maintenance of the proper nuclear position, morphology ^8^ and mediation of various signal transduction pathways between the cell surface and the nucleus ^9^.

SYNE2 mutations were first reported to be associated with autosomal dominant Emery-Dreifuss muscular dystrophy (EDMD) after the discovery of the nesprin-2 T89M mutation, which disrupts the LINC complex and leads to nuclear morphological changes ^10^. EDMD is known to be caused by mutations in nuclear envelope proteins, such as emerin and lamin A/C^11^. Expression of nesprin isoforms is highly tissue-dependent; the short (N-terminally deleted) nesprin-1α2 and nesprin-2α1 isoforms, which lack the CH domain ^5^, are found almost exclusively in cardiac and skeletal muscle ^12, 13^. A double knockout of SYNE1/2 in mouse cardiomyocytes resulted in altered cardiac nuclear position and shape, mislocalization of the LINC complex, altered nuclear biomechanical gene response, and early onset cardiomyopathy^14^. In addition, deletion of nesprin-1 C-terminal KASH domain in mice results in cardiomyopathy associated with cardiac conduction system defects ^15^, indicating that short nesprin isoforms may play vital roles in cardiomyocyte function.

Here we investigate the regulatory activity of two linked AF associated SNPs on the expression of SYNE2 mRNA and protein isoforms in human LAA. We found that the risk alleles of these SNPs decreased expression of the short nesprin-2α1 isoform, which may work as dominant-negative inhibitors by competing with the nesprin-2 giant isoform. We determined that either KD of all SYNE2 isoforms or overexpression of SYNE2α1 increased the nuclear size and decreased nuclear stiffness in iCMs. We hypothesize that the non-risk reference allele may protect the beating atrial cardiomyocyte nucleus from the stress of repetitive motion, and thus protect against AF.

## METHODS

### Human Left Atrial Tissue Processing, RNAseq, and gene-based eQTL analysis

Human LAA tissues were obtained from patients undergoing elective surgery to treat AF, valve disease, or other cardiac disorders, and from nonfailing donor hearts not used for transplant. The living donors all provided informed consent, and consent was obtained from family members for the heart transplant donors, under protocols approved by the Cleveland Clinic IRB. Sample handling, the demographics of these subjects, RNAseq by paired-end 150bp sequencing, and gene-based eQTL analyses were as previously described in Hsu et al. ^4^.

### Transcript isoform-specific eQTL analysis

Quantification of transcripts was done using Kallisto ^16^. QTLTools ^17^ was used to run genome-wide cis-associations (1 MB windows). 1000 sample permutations were used to estimate false discovery rates. The mbd mode of QTLTools was used to verify the identity between genotypes and RNA-seq samples.

### Quantitative Reverse Transcriptase-polymerase Chain Reaction (qRT-PCR)

RNA isolation and cDNA preparation were performed as previously described ^18^. To prepare the 20 μl master mix for each sample, 10 μl of the TaqMan gene expression master mix (Applied Biosystems) was used along with 0.36 μl of 50 μM custom-designed SYNE2 short and long isoform primer and 0.5 μl of 10 μM probe set (Table 1, obtained from Sigma-Aldrich) and 1 μl of the primer limited cyclophilin A (PPIA) primer/probe mix (assay number Hs04194521_s1 from Applied Biosystems). 50 ng of cDNA was added into each well of the 96-well plate. qRT-PCR was performed in a Bio-Rad CRX qRT-PCR machine that had been calibrated for our FAM and VIC fluorescent probes. Thermal cycling started at 95°C for 10 minutes, followed by 40 cycles of 95°C for 15 seconds and 60°C for 60 seconds. Delta Ct values for SYNE2α1 and long isoform expression levels were calculated relative to PPIA expression, and the ΔΔCt method was used to compare expression among samples, yielding log2 based expression values, which we adjusted to a linear scale using 2^−ΔΔCt^.

### Protein extraction and Western blot analysis

Human LAA tissue samples or iCMs were lysed in SDS Lysis buffer containing Tris-HCl, EDTA mixed with protease inhibitor cocktail (P8340, Sigma Aldrich) and 1 mM PMSF. Equal amounts of protein homogenates were separated by NuPAGE 4-12% Bis-Tris gels (ThermoFisher), transferred onto PVDF membranes, and probed with antibodies directed against nesprin-2 (1:1000, ab103020, Abcam) or GAPDH (1:10,000, ab8245, Abcam). Membranes were subsequently incubated with horseradish peroxidase-conjugated goat anti-mouse (GAPDH) or goat anti-rabbit (nesprin-2) secondary antibodies. Signals were detected by the HyGLO HRP detection kit (Denville Scientific) and exposed to Amersham Hyperfilm ECL (GE Healthcare Life Science).

### iCMs culture and siRNA and plasmid transfection

iCMs were purchased from Cellular Dynamics International, the following culture and transfection were underwent according to the manufacturer’s protocol. Silencer^®^ Select scramble siRNA (ThermoFisher Scientific, catalog # 4390843) and SYNE2 siRNA (ThermoFisher Scientific, catalog # 4392420; assay ID: s23328, s23329, s23330) were reconstituted in RNase-free water at 10 µM. iCMs were transfected with TransIT-TKO (Mirus) and scramble or SYNE2 siRNA. A GFP-SYNE2α1 short isoform fusion protein expression plasmid driven by the αMHC-short promoter was constructed by Vectorbuilder (Vector ID: VB171003-1072mch) and transfected into the iCMs using ViaFect transfection reagent (Promega). siRNA or plasmid transfected iCMs were kept in a cell culture incubator at 37°C, 5% CO_2_ for 72 hours and then used for further experiments.

### Reporter gene transfection studies

To examine the promoter activity of the SYNE2α1 isoform, we performed PCR on genomic DNA from two subjects homozygous for either the risk or reference allele of rs11525291 using primer: CAGGTACCACCTGGGCTGCTCAGAGTTTACG and CACAAGCTTCATTCTCAGAGGCCTCCTTTTCATCTTC. The resulting 1195 bp fragment spanning from –1108 to +87 relative to the SYNE2α1 transcription start site (TSS) was digested with Kpn1 and Sac1 for cloning into the promoterless firefly luciferase plasmid pXP1^19^. DNA sequencing confirmed that the only difference between these plasmids was rs11525291. To examine enhancer activity, 60-mer ds-oligos were synthesized (Sigma-Aldrich) for both alleles of either rs1152591 or rs1152595, with the SNPs in the center and HindIII overhangs at the ends. These were cloned into the pT81Luc firefly luciferase plasmid driven by 81 bp of the Herpes simplex virus thymidine kinase promoter^19^. 2 μg of each of these reporter plasmids along with 100 ng of pRL-TK (Promega), a Renilla luciferase expression plasmid to control for transfection efficiency, were transfected into iCMs using ViaFect transfection reagent (Promega). 48 hours after transfection, cells were harvested, and luciferase activities were analyzed using the dual-luciferase reporter assay system (E1910, Promega).

### Immunofluorescent microscopy and nuclear area analysis

iCMs seeded on glass-bottom wells were fixed with 10% phosphate-buffered formalin for 15 min at 37°C, then washed and blocked with 0.1% Triton X-100 or saponin in Casein Blocker (ThermoFisher) in PBS for 30 min. Cells were incubated with lamin A/C mouse monoclonal antibody (1:100; Santa Cruz Biotech sc-376248) and rabbit Nesprin-2 polyclonal antibody (1:200; Abcam ab103020) in blocker overnight at 4°C, washed three times with PBS, and then incubated for 60 min at room temperature with Alexa-Fluor 568 or 488 goat anti-mouse or anti-rabbit secondary (1:1000; Invitrogen) in blocker. Cells were washed three times and mounted with DAPI-containing mounting medium. The nuclear area and circularity of DAPI stained cardiomyocytes were determined automatically after image acquisition using a 20x objective lens on the Cytation3 Cell Imaging Multi-Mode Reader (BioTek).

### Atomic force microscopy

An MFP-3D-Bio atomic force microscope (AFM; Oxford Instruments) mounted on an inverted optical microscope (Nikon Eclipse Ti) was used to obtain force-displacement curves from the cells. A polystyrene bead of 4.5 µm was used to modify the tipless cantilevers (Arrow TL1, nominal spring constant-0.03 N/m) to avoid damage to cells, and the actual cantilever spring constant was determined using a thermal calibration menthod^20^. SYNE2 siRNA and scr siRNA transfected iCMs, SYNE2α1-GFP+ and GFP− iCMs were maintained at 37 °C in the maintenance medium using a custom petri-dish throughout the live-cell nanoindentation assay. For each experiment, at least 40 cells were randomly selected and force curves were obtained on the nuclei at approach/retraction velocity of 5 µm/s. The Young’s modulus was determined from the force-indentation curves using a Hertz’s contact model:

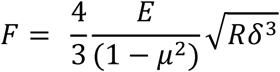

where *F* is indentation force, *E* is Young’s modulus, *µ* is Poisson’s ratio (0.5 for cells), *R* is tip radius (2.25 µm) and *δ* is indentation depth (~500 nm).

### Statistical analysis

GraphPad Prism was used to assess normality and simple statistical analyses. Statistical significances for normally distributed data were assessed by ANOVA followed by Newman-Keuls Multiple Comparison Test (3 conditions) or two-tailed unpaired t-tests (2 conditions). Statistical significances for non-normally distributed data were assessed by Kruskal-Wallis nonparametric ANOVA followed by Dunn’s Multiple Comparison Test (3 conditions) or Mann-Whitney nonparametric two-tailed unpaired t-tests (2 conditions). Differences between conditions were considered significant at p<0.05. All data are shown as mean ± SD unless specified otherwise.

## RESULTS

### AF associated rs1152591 and rs1152595 are eQTLs for SYNE2α1, a short mRNA isoform

235 LAA tissues were obtained during cardiac surgery from subjects of European ancestry and 30 self-reported African Americans, the clinical characteristics were reported in our previous study^4^. In that study, we performed a gene-level transcriptome analysis and we found that the AF GWAS SNP rs1152591 was a strong cis-eQTL for SYNE2 gene expression (q=6.7E-17) ^4^. Here we performed a transcript isoform-specific cis-eQTL analysis for SYNE2 expression. The SYNE2 giant isoform (ENST00000357395.7, called long isoform here) has 116 exons, and the position of the GWAS SNP is shown in Figure 1A. We identified the most abundant LAA SYNE2 mRNA transcript as a short isoform (ENST00000458046.6, referred to hereafter as SYNE2α1 isoform) that initiates transcription at exon 108 of the full-length transcript (Figure 1 A), leading to an N-terminal deleted 62 kDa protein called nesprin-2α1. The AF GWAS SNP rs1152591, is located only 11 bp upstream of the SYNE2α1 isoform transcription start site (TSS) in the intron between exons 107 and 108 (Figure 1A). The GWAS SNP, rs1152591, was a significant eQTL for SYNE2α1 expression (Figure 1B). However, the strongest cis-eQTL was identified as rs1152595, located 6,128 bp upstream of the SYNE2α1 TSS between exons 102 and 103, which is in linkage disequilibrium (LD) with the GWAS SNP (r^2^= 0.78. D’=1) (Figure 1B). Our LAA RNAseq demonstrated abundant expression of SYNE2α1 and ~1000-fold lower expression of SYNE2 long isoform (Figure 1C). However, there is most likely poor capture of the long SYNE2 mRNA by RNAseq, as qPCR specific for the long isoform (exons 8-9) and SYNE2α1 (specific for the unique transcribed sequence in exon 108) only demonstrated ~12.5-fold higher expression of the short SYNE2α1 isoform (Figure 1D).

**Figure 1.**
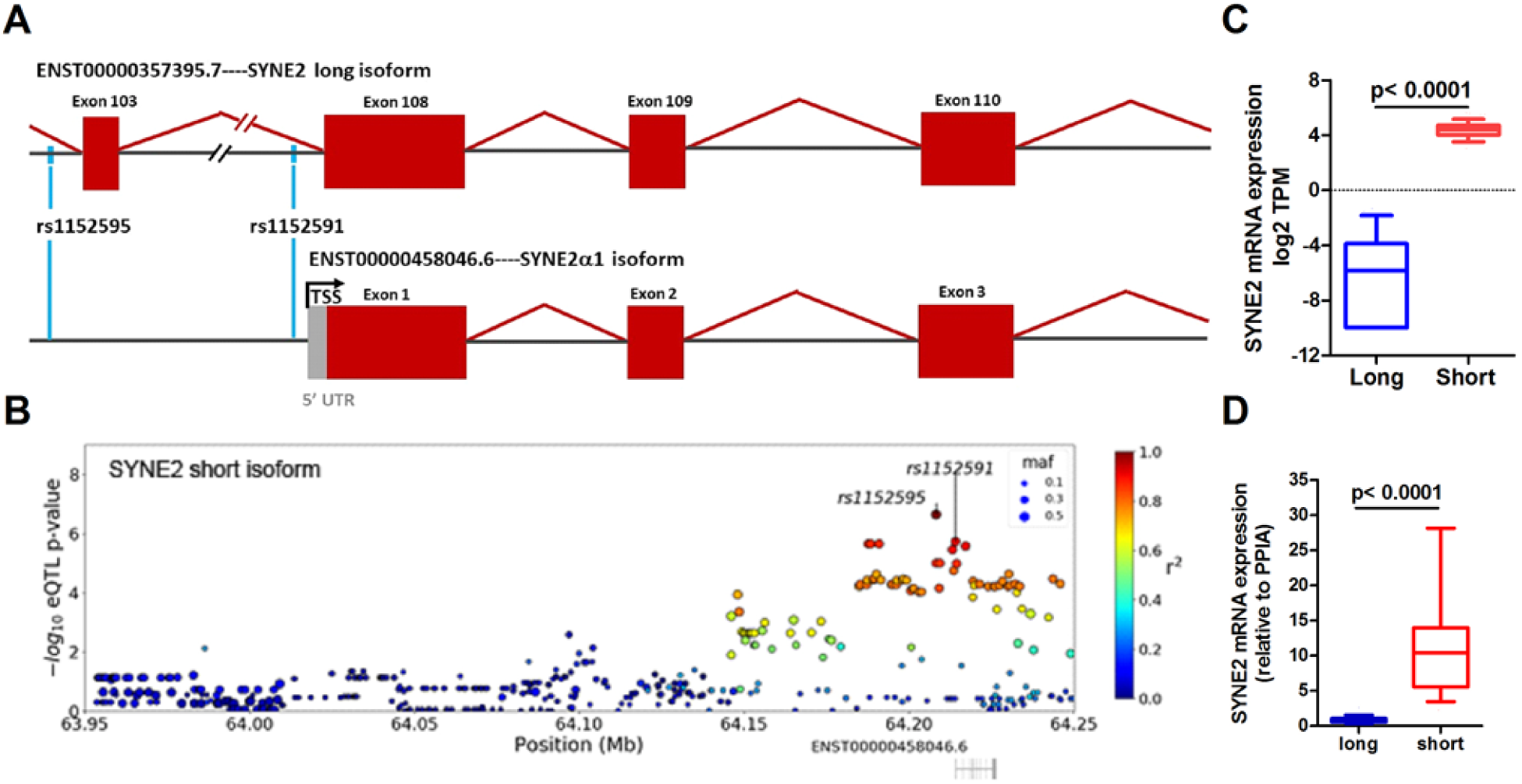
AF risk alleles of SNPs rs1152591 and rs1152595 are associated with decreased expression of SYNE2α1 in LAA. **A**. Schematic diagram of SYNE2 long and SYNE2α1 transcript isoforms (not to scale). **B**. eQTL Manhattan plots of the –log10 p-values for the association of each cis-SNP with the expression of SYNE2α1 mRNA isoform showing rs1152595 as the top eQTL SNP, and the GWAS SNP rs1152591 as the second top SNP. The color bar shows linkage disequilibrium with top cis-eQTL SNP. **C.** LAA RNAseq showed SYNE2α1 (short, red box and whiskers showing median, interquartile range, 10^th^ to 90^th^ percentiles) expression was ~1000 fold higher than the SYNE2 long isoform (blue box and whiskers). **D**. qPCR in 31 LAA RNA samples demonstrated SYNE2α1 expression was 12.5 fold higher than the SYNE2 long isoform.

### Risk alleles of rs1152591and rs1152595 are associated with decreased SYNE2 short isoform expression in LAA tissues

To determine how the risk alleles of rs1152591 and rs1152595 are associated with the expression of SYNE2 isoforms in human LAA tissues, we separated the LAA subjects by genotypes at these two SNPs and found that the AF risk alleles of both SNPs were associated with lower expression of SYNE2α1 (linear trend test p<0.0001), but not the long isoform (Figure 2A). We confirmed this genotype effect on SYNE2α1 expression by qPCR of LAA RNA from 31 subjects, those that carried 1 or 2 vs. 0 risk alleles had lower SYNE2α1 expression (Figure 2B), and the lack of difference between carriers of 1 vs. 2 risk alleles may simply be due to low sample size.

**Figure 2.**
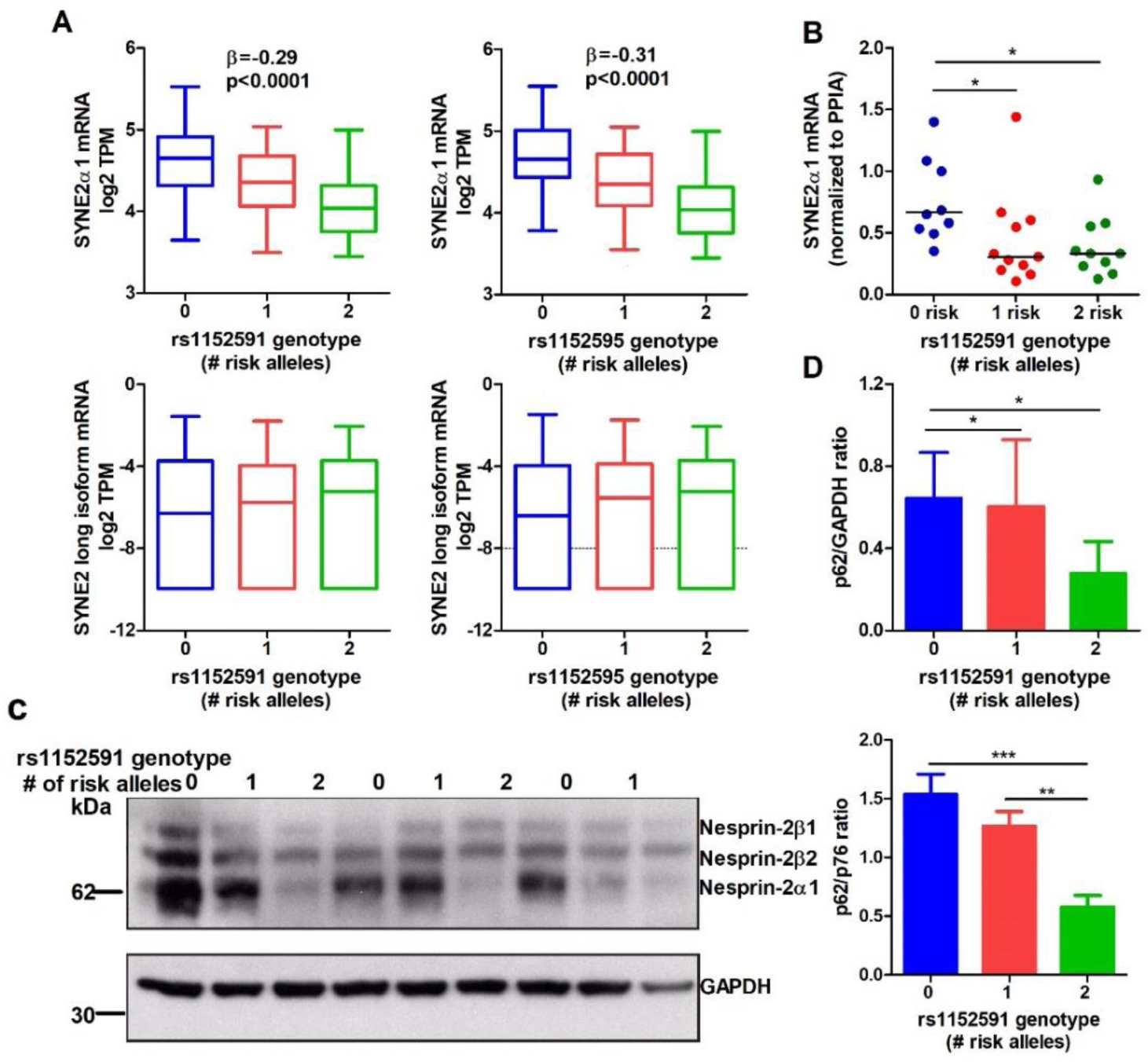
AF risk alleles of rs1152591 and rs1152595 are associated with decreased expression of nesprin-2α1 isoform in human LAA. **A**. Partial regression plot showing the genotype dosage effects of rs1152591 (top GWAS SNP) and rs1152595 (top eQTL SNP) on the expression of the SYNE2α1 and long isoform in LAA from 230 European descent subject. Blue boxes are subjects homozygous for the reference allele of rs1152591 or rs1152595, green boxes are heterozygotes, and red boxes are homozygous for the risk alleles, respectively (boxes show median, interquartile range, and 10^th^ – 90^th^ percentiles). **B**. qPCR of 31 AF LAA RNA samples showing the risk allele of rs1152591 is associated with lower SYNE2α1 mRNA expression (n=31 subjects). **C.** Nesprin-2 western blot of LAA lysates. **D.** Nesprin-2α1 (62 kDa) expression normalized by either GAPDH or nesprin-2β2 (76 kDa) isoform (n=18 subjects), showing subjects with 1 or 2 AF risk alleles have significantly decreased expression of nesprin-2α1 isoform (*, p<0.05; **, p<0.01; ***, p<0.001).

In addition, we performed western blot on 18 LAA lysates using an anti-C terminal nesprin-2 antibody that reacts with the full length and N-terminal deleted short nesprin-2 isoforms. In accordance with the previous reports ^5, 21^, we observed three bands: the band of 62 kDa represents nesprin-2α1, which is only detectable in skeletal, cardiac and smooth muscle; the other two bands are nesprin-2β2 (76kDa) and nesprin-2β1 (87kDa) (Figure 2C). Samples with more risk alleles expressed significantly less nesprin-2α1 when normalized to nesprin-2 beta2 (p76) or GAPDH (Figure 2C, D).

### Risk alleles of rs1152591 and rs1152595 have decreased enhancer activity

The GWAS SNP, rs1152591, and to a lesser extent the top eQTL SNP, rs1152595, were found centered over DNAase I hypersensitivity sites in the fetal human heart (Figure 3A, B). As rs1152591 is 11 bp upstream the SYNE2α1 mRNA TSS, we investigated the effect of this SNP on the strength of the SYNE2α1 promoter. We performed reporter gene transfections using 1196 bp promoters of the risk and reference alleles driving firefly luciferase, which was co-transfected into iCMs along with a Renilla luciferase plasmid used to control for transfection efficiency. Compared to the promoterless plasmid, the risk allele increased reporter gene expression by 7.5-fold, while the reference allele increased expression by 25.8-fold (Figure 3C, all values different from each other, p<0.001). To exam just the proximal region of rs1152591 for enhancer activity, we inserted a 60 bp double-stranded oligos of the two alleles of into a firefly luciferase plasmid driven by a minimal viral promoter. Compared to the enhancerless plasmid, the risk allele of rs1152591 had 2.0-fold enhancer activity, while the reference allele had 4.0-fold enhancer activity (Figure 3D, all values different from each other, p<0.001). The eQTL SNP, rs1152595 was also tested similarly for enhancer activity, and compared to the enhancerless plasmid, the risk allele had no significant enhancer activity, while the reference allele had 2.7-fold enhancer activity (Figure 3E, reference allele different from other two, p<0.001). Thus, both of these linked SNPs were regulatory variants that may alter SYNE2α1 expression, consistent with the risk alleles of both SNPs leading to lower expression of SYNE2α1 mRNA and lower nesprin-2α1 isoform in LAA tissue.

**Figure 3.**
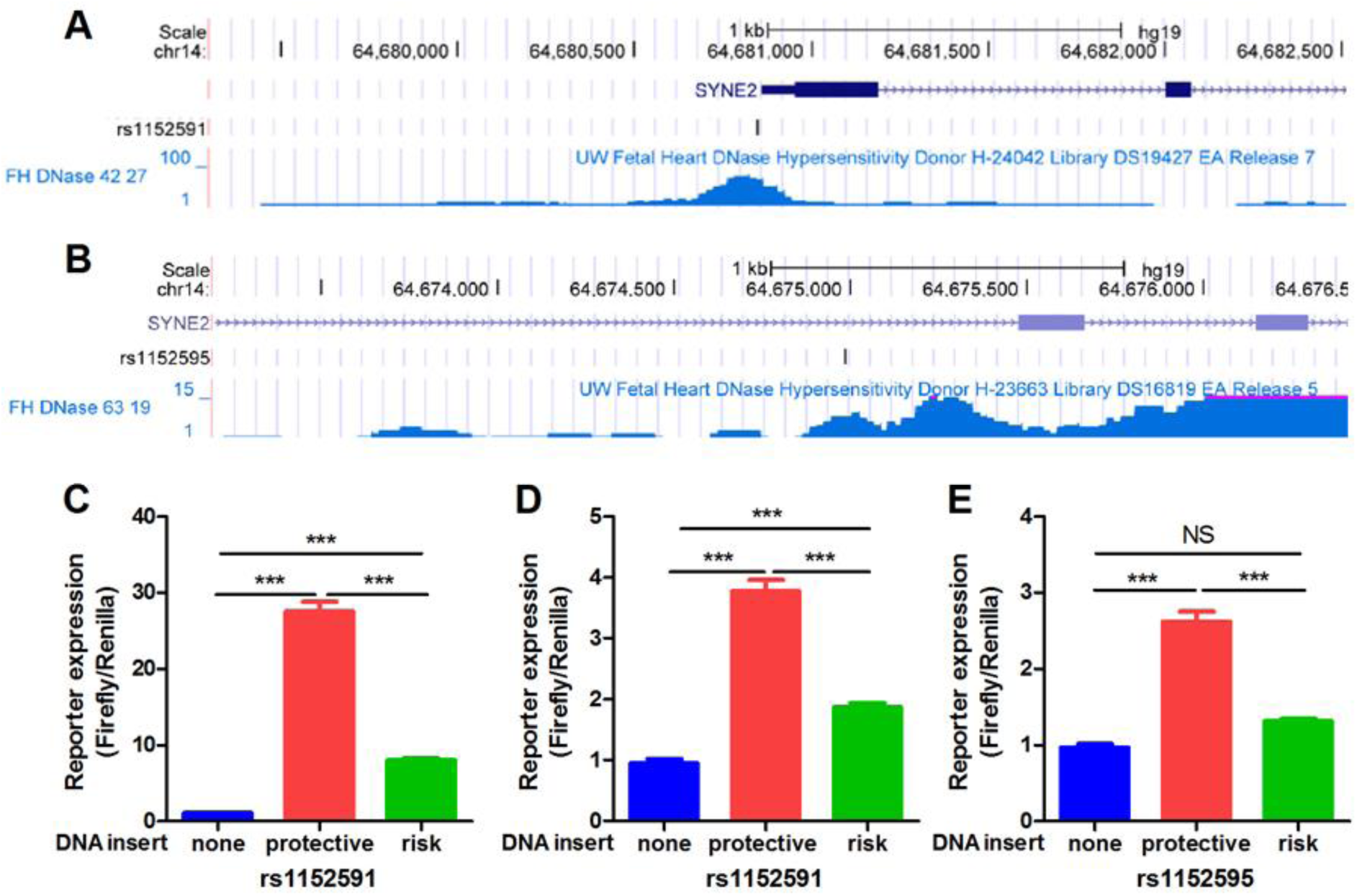
Risk alleles of rs1152591 and rs1152595 have lower promoter and enhancer activity than the reference allele. **A, B.** Browser view of fetal heart DNAase I hypersensitivity of rs1152591 and rs1152595 from ENCODE. **C.** 1.2 kb promoter region of the SYNE2α1 isoform with the risk or reference allele of rs1152591 tested for promoter activity by reporter gene transfection. **D, E.** Enhancer activities of 60-mer oligos containing reference or risk alleles of rs1152591 (D) or rs1152595 (E) in reporter gene transfections. Representative of three independent experiments with n≥3 wells per transfection (***, p<0.001 by ANOVA with Newmann-Keuls posttest).

### SYNE2 KD and short isoform overexpression lead to enlarged iCMs nuclear area

To investigate the effects of altered nesprin-2α1 expression on cardiomyocyte nuclei, we designed an α MHC-driven GFP-SYNE2α1 fusion protein expression vector. 72 h after transfection into iCMs, we measured the nuclear area in GFP+ (overexpressing the nesprin-2α1 fusion protein) and GFP-(non-transfected) cells (Figure 4A). Compared with GFP− control cardiomyocytes, those overexpress nesprin-2α1 isoform had 12.5% larger nuclei (p<0.001, Figure 4B). We used SYNE2 siRNA to KD all SYNE2 isoforms. Western blot and qPCR assays showed a significant reduction of nesprin-2α1 as well as other isoforms 72 h after siRNA transfection (Figure 4C, D), and nuclear measurement revealed that SYNE2 KD cells have 12.6% larger nuclear size (p< 0.001, Figure 4E). Thus, increased nesprin-2α1 expression behaves like a dominant-negative and has the same effect on the nuclear area as KD of all nesprin-2 isoforms.

**Figure 4.**
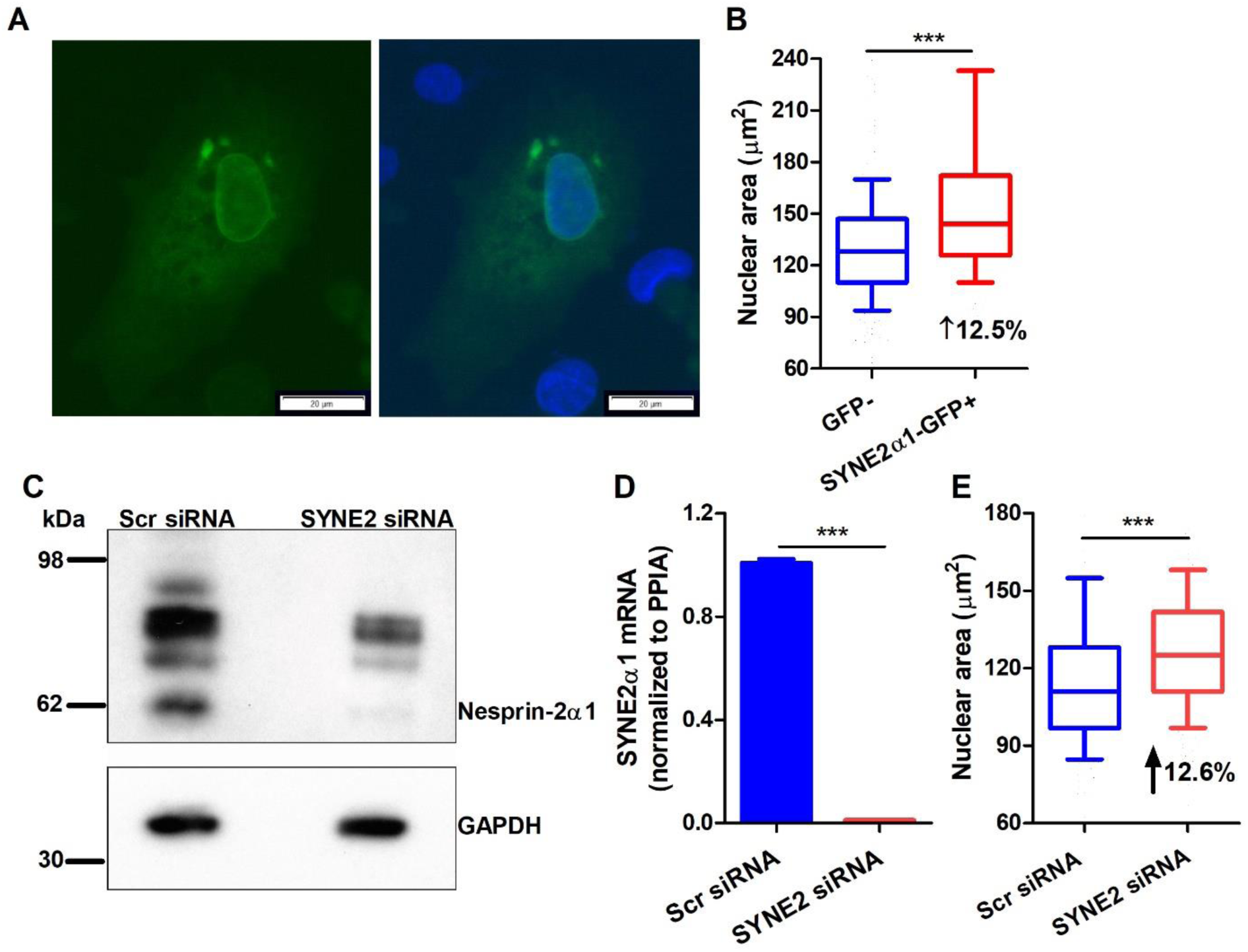
SYNE2α1 overexpression or KD of all SYNE2 isoforms increases the nuclear area in iCMs. **A**. Transient transfection of GFP−SYNE2α1 plasmid into iCMs yielded GFP expression concentrated in the nuclear envelope (left side GFP only, right side GFP and dapi stained nuclei; scale bar = 20 μm). **B**. GFP positive cells overexpressing nesprin-2α1 isoform had a 12.5% increase in median nuclear area vs. adjacent GFP negative cells. **C, D**. Western blot and qPCR showed SYNE2 siRNA KD remarkably lowered several nesprin-2 isoforms, and strongly decrease nesprin-2α1 isoform (62 kDa) and SYNE2 mRNA expression. **E**. Compared with scr siRNA cells, all SYNE2 isoforms KD in iCMs led to a 12.6% increase in median nuclear area. B and E, *** p<0.001 by Mann Whitney two-tailed t-test; D, ***, p<0.001 by two tailed t-test..

### SYNE2 KD and SYNE2α1 overexpression change nuclear biomechanics

Nesprin-2 is a vital structural protein in the nuclear membrane, and our data showed that knockdown of all nesprin-2 isoforms or overexpression of nesprin-2α1 led to larger nuclei in iCMs. To determine effects on nuclear mechanics, atomic force microscopy was performed to assess the nuclear stiffness, using Young’s modulus as a measure of nuclear stiffness. We compared SYNE2α1-GFP+ and GFP− cells within the same dish and found that overexpression of the nesprin-2α1-GFP fusion protein led to a 33% decrease in nuclear stiffness (from 3.17 to 2.11 kPa, p<0.0001, Figure 5A). Similarly, comparing SYNE2 siRNA vs. scr siRNA, KD of all SYNE2 isoforms led to a 57% decrease in nuclear stiffness (from 3.09 to 1.31, p< 0.0001, Figure 5B). Representative force curves for the indentation of nuclei from control, nesprin-2α1 overexpressing, and SYNE2 KD iCMs are shown in Figure 5C, again confirming that the SYNE2α1 isoform behaves as a dominant-negative, reducing the stiffness of the nucleus as KD of all SYNE2 isoforms. Lamin A/C immunofluorescent staining was performed to determine if SYNE2 KD altered the lamin A/C structure or localization in the inner nuclear membrane; and, no differences were observed (Figure 5D). Thus, SYNE2 effects on nuclear stiffness were not due to changes in the lamin A/C nuclear lamina.

**Figure 5:**
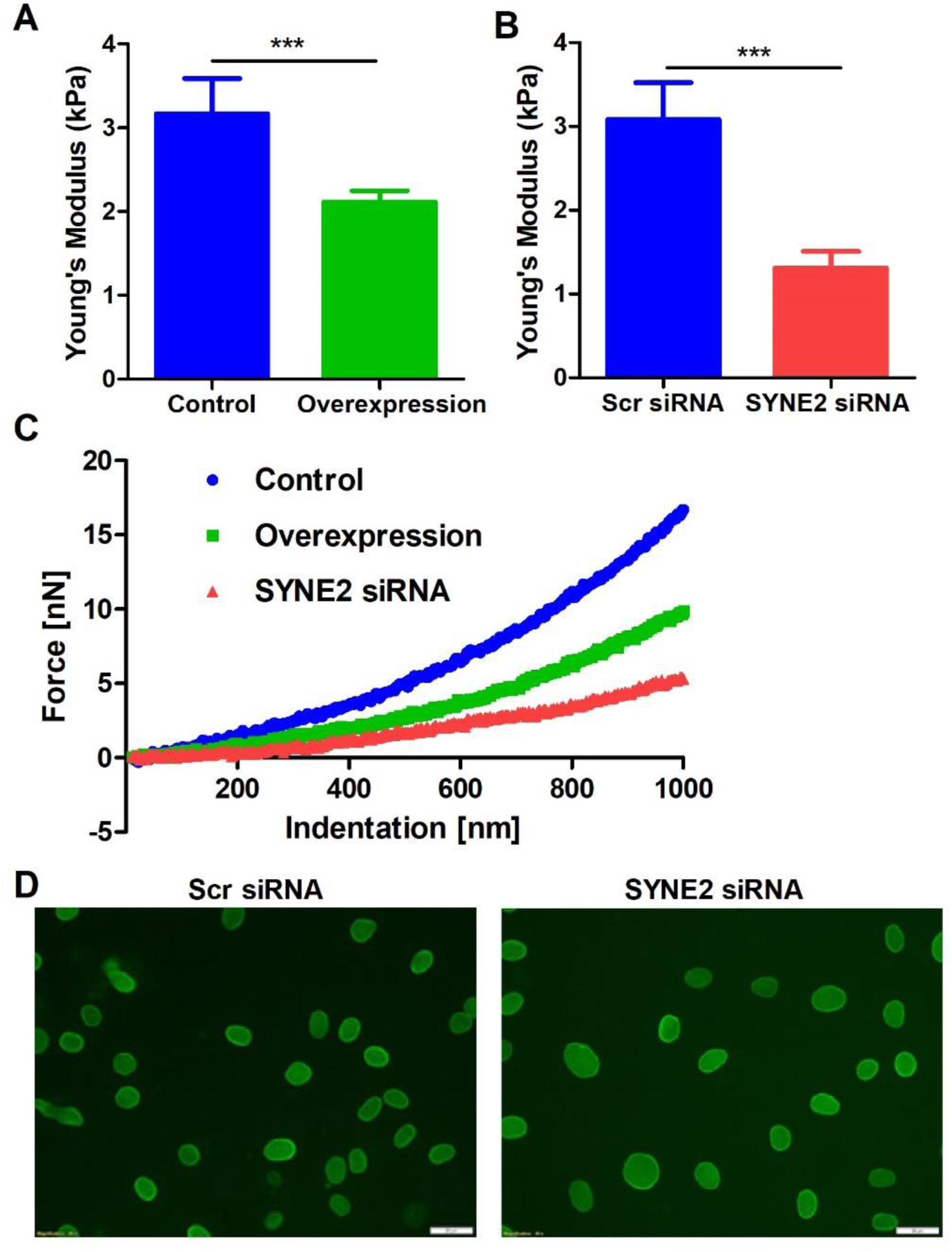
SYNE2α1 overexpression or KD of all SYNE2 isoforms decreases nuclear stiffness in iCMs. **A, B**. Young modulus values for control and SYNE2α1−GFP+ cells, scr siRNA and SYNE2 siRNA KD cells 72h after transfection (mean ± SD, ***, p<0.0001 by two-tailed t-test, number of cells: GFP−, n=60; SYNE2α1−GFP+, n=60; scr siRNA, n=40; SYNE2 siRNA, n=60). **C**. Representative AFM force curves for GFP−, SYNE2α1−GFP+, and SYNE2 KD cells, showing the force needed to push varying depths into the nucleus. **D.** Lamin A/C immunostaining in scr siRNA and SYNE2 siRNA KD iCMs (scale bar = 20 μm).

## Discussion

We found that AF SNPs rs1152591 and rs1152595 were gene dosage-dependent eQTLs for the SYNE2α1 mRNA isoform encoding nesprin-2α1, with the risk alleles associated with decreased expression of nesprin-2α1 in human LAA tissue. Nesprin-2 is an important LINC complex protein, which links the nucleus to the cytoskeleton ^5–7^. By physically connecting the nucleus with the cytoskeleton, the LINC complex participates in many cellular activities including maintaining nuclear morphology, regulating nuclear position, and mediating mechanical cell signaling and downstream gene regulation ^22–24^. Previous nesprin-2 knockout mouse models showed increased nuclear area in cardiomyocytes ^14^ and fibroblasts^25^. Nesprin-1 and nesprin-2 double knockout mice have early onset cardiomyopathy ^14^. In addition, truncation of the N-terminal nesprin-2 CH domain in mice led to epidermal thickening increased epidermal nuclear size, and altered nuclear shape ^26^, demonstrating dominant-negative activity. Overexpression of the C-terminal KASH domain resulted in the displacement of nesprin-2 giant, disruption of the integrity of LINC complex, and decreased connection of the cytoskeleton from the nucleus ^27^. Our engineered iCMs confirmed that nesprin-2α1 acts as a dominant-negative, that SYNE2 KD or nesprin-2α1 overexpression in iCMs give rise to the similar phenotypes such as decreased nuclear stiffness and larger nuclear area, without any apparent influence on lamin A/C integrity (Figure 6).

**Figure 6:**
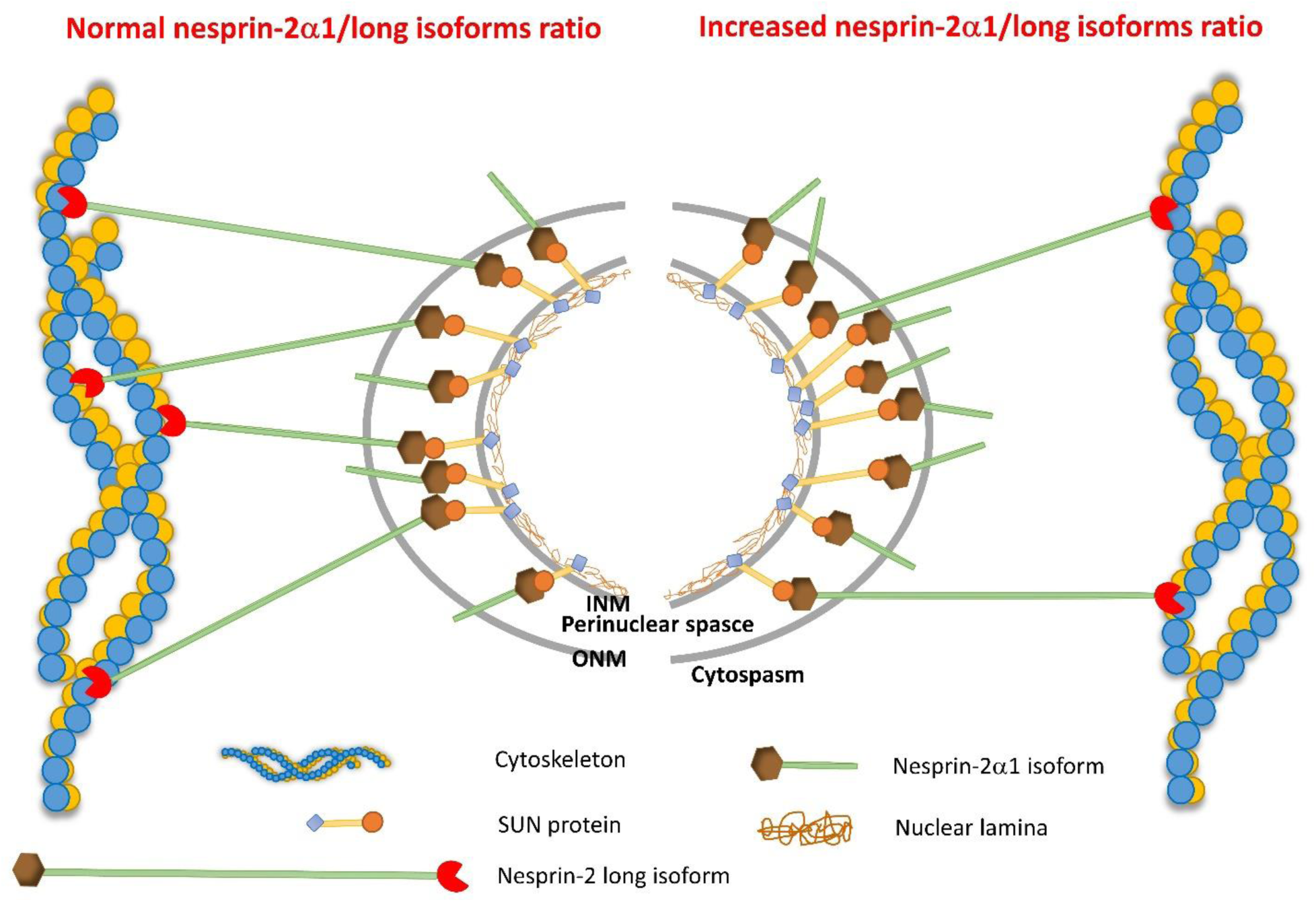
Model for the dominant-negative activity of nesprin-2α1. KD of all SYNE2 isoforms reduces the levels of nesprin-2 giant, disconnecting the nuclear membrane from the cytoskeleton, allowing the nucleus to enlarge and decreasing nuclear stiffness. Similarly, nesprin-2α1 overexpression competes at the nuclear membrane with the nesprin-2 giant, but it cannot bind to the actin cytoskeleton, thus it acts as a phenocopy of SYNE2 KD with dominant-negative activity.

Our previous eQTL analysis indicated that rs1152591, the top AF GWAS SNP at this locus, was a strong eQTL for SYNE2 expression at the gene level (p=6.13E-17) ^4^. Here we performed transcript-level analysis, and we found that the linked SNPs rs1152595 and rs1152591 were the top two significant eQTLs for the SYNE2α1 isoform, and our reporter gene transfection showed that the risk alleles of both SNPs decreased enhancer and/or promoter activities, suggesting that these two eQTL SNPs regulate SYNE2α1 expression, and thereby play a role in the AF susceptibility. We further demonstrated that the rs1152591 risk alleles were associated with lower SYNE2α1 and nesprin-2α1 expression in LAA samples.

Lamin A/C is the major contributor to nuclear stiffness, with loss of lamin A/C leading to more deformable nuclei and metabolic consequences in mice ^28^. In contrast, increased lamin A/C expression, or specific LMNA mutations such as D192G, E161K or N195K, cause stiffer, less deformable nuclei and lead to dilated cardiomyopathy ^29–31^. Our vitro studies showed consistent result that both nesprin-2 KD and nesprin-2α1 overexpression decrease nuclear stiffness in the cardiomyocytes that indicate the cardiomyocytes with risk alleles tend to have stiffer nuclei. Similarly, the nuclei of HGPS cells are stiffer and have more sensitive to mechanical stress than normal cells ^32^. The proper structure/function of the lamin network protects cells from mechanical stretching induced nuclear rupture and DNA damage ^33^, influences the posttranslational modification of histones (e.g., acetylation and methylation) as well as the dynamics and intranuclear localization of heterochromatin ^34^, and influences multiple features of cell behavior including motility^35^, polarity, and cell survival ^36^. In a similar fashion, the AF risk alleles, leading to low expression of nesprin-2α1 in cardiomyocytes, may strengthen the connection of the cytoskeleton to the nuclear membrane and lamin network. This enhanced cytoskeletal-nuclear connection would increase nuclear stiffness and the response to constant mechanical stress during contraction, leading to more nuclear damage and cell death that may promote AF susceptibility.

## Nonstandard Abbreviations and Acronyms

AF: Atrial fibrillation
CH: Calponin-homology
cis-eQTL: cis-expression quantitative trait locus
EDMD: Emery-Dreifuss muscular dystrophy
iCMs: Human stem cell-derived induced cardiomyocytes
KASH: Klarsicht-ANC-Syne-homology
KD: Knockdown
LAA: Left atrial appendage
LD: Linkage disequilibrium
LINC: The linker of nucleoskeleton and cytoskeleton
SR: Spectrin-repeat
Scr: Scramble
SUN: Sad1p/UNC-84
TSS: Transcription start site

